# Consciousness as a Concrete Physical Phenomenon

**DOI:** 10.1101/557561

**Authors:** Jussi Jylkkä, Henry Railo

## Abstract

The typical empirical approach to studying consciousness holds that we can only observe the neural correlates of experiences, not the experiences themselves. In this paper we argue, in contrast, that experiences are concrete physical phenomena that can causally interact with other phenomena, including observers. Hence, experiences can be observed and scientifically modelled. We propose that the epistemic gap between an experience and a scientific model of its neural mechanisms stems from the fact that the model is merely a theoretical construct based on observations, and distinct from the concrete phenomenon it models, namely the experience itself. In this sense, there is a gap between any natural phenomenon and its scientific model. On this approach, a neuroscientific theory of the constitutive mechanisms of an experience is literally a model of the subjective experience itself. We argue that this metatheoretical framework provides a solid basis for the empirical study of consciousness.

## 1. Introduction

The current scientific view about the relationship between consciousness and neural activity is ambivalent. On the one hand, the enterprise to solve the mystery of consciousness using neuroscience is built on the physicalistic idea that consciousness is identical with neural activity: We know that the brain is necessary for consciousness, and neuroscience has begun to unravel the neural mechanisms that enable subjective, conscious experience (Dehaene & Changeux, 2011; Koch, Massimini, Boly, & Tononi, 2016; Laureys, 2005). On the other hand, scientists and philosophers often assume that science can merely observe *correlations* between neural activity and consciousness, not consciousness per se. After all, how could neural activity be *identical* with consciousness given that the two appear so different? For this reason, neuroscientists search for the neural correlate of consciousness (NCC), which has been defined as the minimal set of neural processes that are together sufficient for a specific conscious experience (Crick and Koch, 1990). Here, we attempt to shed light on this discrepancy.

The present article is an attempt to formulate a plausible and solid metatheoretical foundation for the neuroscientific study of consciousness. This is, in our view, an imperative objective for consciousness science. While consciousness science has established itself as a specific field of empirical science, it seems to be still searching for its identity: Arguably, it is a field that does not yet have a clearly defined explanandum. This is likely to foster false disagreements between theories of consciousness, and contribute to the perception that consciousness science is a less rigorous field than other fields of science (Michel et al., 2018).

In this paper, we defend a form of physicalistic monism, which holds that all concrete phenomena are physical—this is typically the starting point in modern science. Complex phenomena, such as those studied by chemistry, biology, or neuroscience, belong to the same ontological category as fundamental physical processes (those studied by fields such as particle physics), and are based on them. We could say that “physical” denotes the set that contains all concrete phenomena, and terms like “biological” or “neural” denote subsets of that all-encompassing set. By the term “physical” we mean the ontological category of phenomena studied by physics—thus, on our reading “physical” is a *placeholder* for whatever is the fundamental nature of phenomena studied by physics. As we will argue below, we take subjective experiences or consciousness to be concrete natural phenomena (Revonsuo, 2006; Searle, 2002; Strawson, 2008b). If we endorse the physicalistic premise that all concrete natural phenomena are physical, it follows that our experiences are physical phenomena. This might appear to be a contradictory claim, given that the term “physical” is often defined in opposition to “mental”. For instance, according to the Oxford Dictionary, “physical” is something “relating to the body *as opposed to the mind”*, or as “relating to things perceived through the senses *as opposed to the mind”* (emphasis added). Here we do not adhere to this standard usage of “physical”, because it *a priori* commits us to dualism. On our reading, terms like “experiental” or “conscious” denote certain natural phenomena which belong to the class of physical phenomena.

The challenge for physicalists who consider experiences to be identical with some neural processes is to explain why the two *appear* so different, or why there is an epistemic gap between the two. For instance, a person with total congenital achromatopsia arguably cannot know *what it is like* to see the vivid colors of a sunset on a clear evening, even if she perfectly knew all the details of how the human brain works. It appears that science cannot tell us anything about the qualitative or subjective aspects of experiences—the colors of the sunset, the bitterness of coffee, or the pain of having a manuscript rejected—and thus cannot completely capture their nature (e.g. Chalmers, 1996; Jackson, 1986; Nagel, 1979). The difficulty of conceiving subjective experiences—or “qualia”, to use the philosophical term referring to the “qualitative feelings” associated with experiences—as an object of scientific study has led some empiricists to explicitly avoid referring to them (Dehaene, 2014), or to claim that such aspects do not exist (Dennet, 1991; Dennett, 2018).

Many attempts in philosophy have been made to explain where the epistemic gap stems from, and the so-called phenomenal concepts strategy is probably the most popular alternative (for a good overview, see Balog, 2012). It aims to reduce the gap into differences between how phenomenal and non-phenomenal concepts refer. Non-phenomenal concepts have a mode of presentation that is distinct from their referent (e.g., we recognize water based on its perceptual features, but not everything that appears to be water is water), but phenomenal concepts are suggested to be more intimately tied with their referent. For instance, it is claimed that phenomenal concepts are recognitional concepts whose mode of presentation is identical with their referent (e.g. Loar, 1990); that phenomenal concepts are similar to indexicals like “here” and “now” (e.g. Ismael, 1999); or that they are quotational concepts, so that e.g. “pain” refers to “that state: ___”, where the blanks are filled in with pain itself (e.g. Papineau, 2002). Our main focus in this paper is not on differences between how phenomenal and scientific concepts refer, but instead on the question of how, and to what extent, science can model consciousness.

In this paper we take it as our premise that experiences, in all their qualitative richness, are concrete physical phenomena (Strawson, 2008a). To emphasize that empirical explanations of consciousness aim to characterize the causal physical processes that are identical with consciousness—not processes that merely *correlate* with it—we use the concept “constitutive mechanisms of consciousness” (CMC; Revonsuo, 2006). As discussed later in detail, by CMC we mean all those physical processes that together make up consciousness. Similar to biologists who try to describe the hierarchical physical processes that constitute, for example, a living cell or an amoeba, consciousness science aims to describe the hierarchical constitution of the processes that constitute consciousness. This approach is superficially similar to the aim of describing the NCC. The NCC and CMC approach share the idea that consciousness can be described as a hierarchical process: lower level phenomena (molecules, neurons, their activity) combine to produce a system-level phenomenon (a network of neurons and brain areas), which corresponds to consciousness. The crucial difference between these two views is that the CMC approach implies that the physical phenomenon the scientific theories describe *is* consciousness. The NCC approach, in contrast, suggests that the theories describe a physical phenomenon that merely *correlates* with consciousness. This brings us back to the aim of the present paper: What does it mean to claim that certain neural activation patterns are identical with experiences, given that the two appear so different?

As a summary, we aim to characterize how subjective experiences are related to empirical observations and models about their constitutive mechanisms, and why there appears to be an epistemic gap between the two. We propose that the epistemic gap is distinctness between the scientific CMC-model and the concrete experience. A scientific model is always distinct from the concrete phenomenon it models, and in this sense there is an epistemic gap between any phenomenon and its model, not just experiences. As suggested by Hawking and Mlodinow (2010), through science we can never know the nature of the world in itself, we can only model it based on observations. This is because scientific models (like our everyday models of the world) are phenomena in the scientists’ brains: they are mental models that aim to explain and predict scientific observations of external phenomena. Thus, scientific models of subjective experiences are themselves conscious phenomena in the minds of scientists. Our framework implies that our subjective experiences are different from other natural phenomena solely because they are the only phenomena in the universe that constitute *our* subjective realm. If we assume that all the properties of experiences (including what they feel like) are physical and causally efficacious, then they can causally interact with measuring devices. Thus, it is possible to observe and model consciousness itself, not just its correlates. We call our approach *Naturalistic Monism*, because it shares some key components with Russellian Monism (RM), but is completely naturalistic: it does not postulate the existence of properties beyond the scope of natural science.

This article is structured as follows. In the first part, we explain how our approach is motivated in particular by Strawson’s (2008a, 2008b) version RM, but also how our view significantly differs from it. In the second part of the article, we describe in what sense scientific theories are distinct from the phenomena they model, and what implications this has for consciousness research. In the third section, we discuss the implications of Naturalistic Monism from the perspective of science of consciousness, and conclude that neuroscience can study consciousness, not just its correlates. Finally, in the fourth section we briefly discuss the philosophical implications of our approach.

## 2. Consciousness as a physical phenomenon

Among philosophers, RM has recently gained popularity as an explanation of how experiences are related to their neural mechanisms (e.g. Alter & Nagasawa, 2012; Chalmers, 2016; Goff, 2017; Montero, 2015; Schneider, 2017). It promises to account for why science appears to have trouble in explaining the subjective aspects of experiences, while maintaining that experiences are nevertheless identical with their neural mechanisms. Naturalistic Monism shares two premises with RM: First, it holds that experiences are concrete natural phenomena; Second, it accepts that science has *in some sense* limited access to their nature. The crucial difference between Naturalistic Monism and RM concerns how the second premise should be interpreted.

Strawson (2008a, 2008b) calls his version of RM “Real Materialism”, because it is both materialistic (or physicalistic; Strawson uses the terms interchangeably) and realistic about experiences. He writes:

> “Realistic physicalists, then, grant that experiential phenomena are real concrete phenomena—for nothing in life is more certain—and that experiential phenomena are therefore physical phenomena. It can sound odd at first to use ‘physical’ to characterize mental phenomena like experiential phenomena, and many philosophers who call themselves materialists or physicalists continue to use the terms of ordinary everyday language, that treat the mental and the physical as opposed categories. It is, however, precisely physicalists (real physicalists) who cannot talk this way, for it is, on their own view, exactly like talking about cows and animals as if they were opposed categories. Why? Because every concrete phenomenon is physical, according to them. So all mental (experiential) phenomena are physical phenomena, according to them; just as all cows are animals.” (Strawson, 2008c)

Naturalistic Monism endorses this premise, as it is formulated in the above quote. The existence of consciousness is the most certain thing in the world: we can doubt the existence of the whole external world, but not our own experiences. If we are physicalists and take it that experiences feel like something, then we must admit that there exist at least some physical phenomena in the universe that feel like something. However, this is where our agreement with Strawson and other Russellians largely ends. For Strawson continues:

> “I am happy to say, along with many other physicalists, that experience is ‘really just neurons firing’ […] But when I say these words I mean something completely different from what many physicalists have apparently meant by them. I certainly don’t mean that all characteristics of what is going on, in the case of experience, can be described by physics and neurophysiology or any non-revolutionary extensions of them. That idea is crazy. It amounts to radical ‘eliminativism’ with respect to experience, and it is not a form of real physicalism at all.” (ibid)

Strawson’s idea is probably the following: if it was possible to characterize the nature of experiences completely in terms of science, no room would be left for phenomenal qualities or what-it-is-likeness, because science appears to say nothing about such properties. Strawson appears to claim that if experiences could be perfectly characterized scientifically, we would be philosophical zombies (Chalmers, 1996). We strongly disagree with this claim: as we will argue, it conflates experiences with their scientific models.

Strawson’s view is based on a metaphysical distinction between extrinsic (roughly: relational, structural, and dispositional) and intrinsic (non-relational, non-structural, and non-dispositional) properties (for a useful discussion of the distinction, see Seager, 2006). On this view, science is only concerned with extrinsic properties—how objects behave, which causal dispositions they have, which relations they stand in, what is their structure, and so on. For instance, in Newtonian physics, force is defined in relation to mass and acceleration. For Russellians, this leaves open the question: what is the *non*-relational, *non*-dispositional, and *non*-structural nature of a phenomenon? Russell (1927) himself argued that this intrinsic nature is ontologically “neutral” (that is, neither mental nor non-mental; see Stubenberg, 2016), but Strawson takes it that the intrinsic nature of all physical phenomena is (proto)mental (see also Goff, 2017). To use the current terminology, Strawson would argue that what an experience feels like is part of the intrinsic (science-transcendent) nature of the experience, but CMC characterizes the extrinsic (scientifically observable) nature of the same phenomenon.

The intrinsic-extrinsic distinction could be criticized in many ways (see e.g. Hiddleston, 2019; Howell, 2015; Kind, 2015), but here we simply note that the postulation of intrinsic properties is metaphysically promiscuous and unscientific. If we suppose that there is an unobservable and causally impotent intrinsic property corresponding to each observable, causally efficacious property, we double the number of physical properties in the universe solely based on armchair reflection. Moreover, as argued by Ellis (2001), among others, there may be no need to postulate any categorical properties to “ground” causal dispositions or relational properties, because all natural phenomena can be considered as essentially causal-dispositional, or processes. For instance, scientifically it makes no sense to speak of the nature of quarks independently of how they interact with other subatomic particles. According to the standard version of quantum chromodynamics, it is impossible for quarks to exist in isolation, so it is nomologically impossible for them to possess a non-relational nature.

Because RM implies that experiences are beyond the scope of science, it renders the scientific study of consciousness impossible in principle: science can only study the extrinsic correlates of consciousness, not consciousness per se. This resembles dualistic notion that “conscious experience involves properties of an individual that are not entailed by the physical properties of that individual, although they may depend lawfully on those properties” (p. 110; Chalmers, 1996). This type of reasoning in part underlies the use of the term “correlate” to describe the neural mechanisms that underlie consciousness. If we could account for the epistemic gap without assuming an *ontological* gap between different types of properties, we should do so.^1^ This is the aim of Naturalistic Monism. It holds that experiences are just like any other physical phenomena in that they can be observed and scientifically modelled. The epistemic gap between experiences and their scientific models does not reflect the existence of any separate types of properties, but instead simply the distinctness between a phenomenon and its scientific model.

## 3. Naturalistic Monism: phenomena and their scientific models

Naturalistic Monism shares with RM the Kantian assumption that science has in some sense limited access to the nature of the phenomena it studies. However, as explained above, whereas RM holds that there exist two distinct types of properties (extrinsic and intrinsic), Naturalistic Monism takes it that there exists only one class of physical phenomena. The limits of science are not due to the existence of ontologically distinct types of properties, but instead to the fact that we can only know the world through observations and scientific models.

Schneider (2017), a proponent of RM, motivates the existence of science-transcendent intrinsic properties by quoting the theoretical physicist Stephen Hawking, who famously asked:

> “What is it that breathes fire into the equations and makes a universe for them to describe? The usual approach of science of constructing a mathematical model cannot answer the questions of why there should be a universe for the model to describe.” (Hawking, 1988, 174).

Strawson (2008b), in turn, quotes the astrophysicist Arthur S. Eddington, who noted that

> “[…] science has nothing as to the intrinsic nature of the atom. The physical atom is, like everything else in physics, a schedule of pointer readings. The schedule is, we agree, attached to some unknown background.” (Eddington, 1929, 259)

But do Hawking and Eddington intend that there would exist some *properties* of physical phenomena completely beyond the scope of science, as Russellians would have it? Or did they simply mean that we cannot *know* the nature of physical phenomena without relying on observation (“pointer readings”) or without utilizing scientific models and equations? The former claim is metaphysical, the latter is purely epistemological. At least Hawking intended to make a purely epistemological claim, which he and Mlodinow formulated more explicitly when they introduced the notion of *Model-Dependent Realism* (MDR) (Hawking & Mlodinow, 2010). According to MDR, we can never know the nature of the world as it is in itself, but instead we can only know it through how it affects our senses. Based on observations, we can formulate models of the world in our minds, but we can never step outside the models and compare them to model-independent reality. We can never be certain if our models are true in a strict sense of the word; we can only evaluate whether they are elegant, can predict observations, or can be used to manipulate phenomena.

Hawking and Mlodinow consider scientific models on par with our non-scientific, everyday models of the world: both are in the mind/brain of the organism and based on constant causal interaction between the organism and its environment. Consciousness itself can be considered as an internal model of the world, which affords us to predict and explain what happens in our surroundings, increasing our chances of survival (Friston, 2010; Hobson & Friston, 2014; Revonsuo, 2006). In this sense, there is only a matter of degree between a person forming an inner representation of the world based on their everyday interaction with it, aided by nothing but naked senses and their body, and a researcher building a scientific model based on more sophisticated, instrumentally aided, and theoretically-driven observations and manipulations of the phenomena. In both cases, the model is a natural phenomenon in the brain of the modeler, and can itself be an object of scientific inquiry (more about this in section 5.1.).

It is a philosophical question whether a model can ever be considered as “true”, but from a purely naturalistic perspective we can judge to what extent an organism’s internal model of the world affords it to interact with its environment efficiently. A mouse has a valid internal model of its surroundings if the model affords the mouse to seek shelter from the approaching cat in the nearest burrow. Likewise, our scientific atomic model can be considered as valid if it affords us to build functioning nuclear reactors. Thus, internal models—irrespectively of whether they are of a scientific or everyday variety—can be minimally considered as valid or even “true” in a *pragmatic* sense. Whether they are true in some stricter sense is a philosophical problem that is beyond the scope of the present paper.

According to Naturalistic Monism, the epistemic gap between a subjective experience and its scientific description is distinctness between two phenomena: the experience in the mind of a participant and the scientific model in the mind of the researcher. Naturalistic Monism implies that *an experience is the concrete phenomenon that a scientific model of its constitutive mechanism describes;* it is what underlies scientific observations of its constitutive mechanisms. Scientists can model experiences, but the participants’ experiences are always distinct from their scientific models, which are in the minds of researchers. Whereas the participant has “immediate” or first-person access to their experiences, the researcher can know them “mediately” through observations and models. The researcher does not have direct access to the participant’s experiences; they can only access the participant’s consciousness indirectly by causally interacting with it (and directly experiencing a model of it in their own consciousness). In this respect, consciousness is an exceptional object of research: it is the only phenomenon that we can at least partly know or experience directly, not only based on observations and models. Thus, in the case of experiences, we can directly compare the model and the concrete phenomenon that is modeled—something we can never do in the case of, say, atoms.

Science can model all the aspects of experiences—including what they feel like—*but only model*. For instance, suppose that subjective pain could be modelled as T-type interaction between neural modules M1 and M2 (or T-interaction, for short). “T-interaction” is just pain’s scientific model, formed in the mind of a scientist based on observations, such as which neural events are correlated with reports of pain, avoidance behavior, certain facial expressions, and so on. Even a scientific realist, who assumes that scientific theories can truthfully capture the nature of natural phenomena, cannot hold that pain is *literally* nothing but “T-interaction”, because “T-interaction” is a theoretical *model* of pain, and distinct from pain itself. Instead, the realist should be interpreted as claiming that pain can be truthfully *modelled* as “T-interaction”. This is compatible with the fact that when the concrete process scientifically modelled as “T-interaction” happens in a subject, the occurrence of the process feels like something for her. Crucially, the subjective feel of pain is nothing distinct from the process described by the model; it is the happening of the concrete process itself. Accordingly, a scientific realist can accept that pain has a qualitative feel; what their realism implies is only that *the feel of pain can be truthfully modelled as “T-interaction”*. In this sense, we can eliminate the theoretical notion of “qualia”, conceived of as a non-dispositional, non-relational, and non-structural property, without claiming that experiences would not feel like anything (in the everyday sense of “feeling like something”) (cf. Dennett, 2018).

A neuroscientific CMC_E_-model of an experience E is never identical with E; it only refers to E. Whereas science can only model experiences through observation, we know our own experiences immediately. My pain is not something outside of me that I observe, it is part of my subjective realm. It constitutes the physical process that is my consciousness, which could be observed and modelled by a scientist as CMC_pain_. To paraphrase Edelman and Tononi (2000), “Unlike any other entity, […] with consciousness *we are what we describe* scientifically” (p. 14, italics in the original). The reason why we easily think that there is an epistemic gap only for experiences is because our experiences constitute us, and thereby we have non-empirical access to their nature—access that is not mediated through observations and models. Naturalistic Monism implies that there is an epistemic gap between *any* concrete phenomenon and its scientific model, due to the distinctness between the model and the phenomenon. As discussed later in the article (section 5.2.), how wide we take the gap to be depends on which philosophy of science we endorse.

Naturalistic Monism implies that what an experience feels like is part of its nature in itself, independently of observations, scientific models, or other theoretical characterizations. In fact, already describing an experience as “feeling like something” is already a conceptualization, and it could be debated whether it adequately or truthfully captures the nature of the experience. Descriptions of consciousness typically depend on the person’s favorite philosophical theory, religion, culture, or level of education. Consciousness could be said to be “neural activation”, “qualitative”, “phenomenal”, “feeling like something”, “non-relational”, “non-structural”, consisting of “qualias”, “soul”, “Brahman”, and so on. All such *definitions* of experiences could be questioned, but that does not amount to questioning the nature of experiences themselves. If you take René who believes in an immaterial soul, David who believes in qualia, and Daniel who denies the existence of qualia, and stick them with a needle, the same kind of natural phenomenon is instantiated in all of them. The concrete phenomenon that occurs in them is of the same type, even if they define it in very different ways—nature does not care about our definitions.^2^

The qualitative feel of experiences is often considered as problematic or even impossible to model scientifically. Naturalistic Monism implies that it is possible to model what experiences feel like—but only model. The discrepancy between what, say, pain feels like for a subject S and how a scientific model characterizes it is simply distinctness between pain and its scientific model. Pain is a natural process happening in S, whereas its scientific model is a highly complex conceptual representation in the minds of scientists. What pain feels like for S is not something over and above the process described by the model; instead, it is the *happening* of the described process in S. When pain happens in me, I can minimally say it feels like “this”, referring to the happening of my pain experience (Ismael, 1999; Papineau, 2002). Consciousness science aims to model such processes, and can in this sense model what experiences feel like. In the form of an argument:

1. What pain feels like for S = the happening of pain in S.
2. The happening of pain in S is a concrete physical process.
3. Science can model all concrete physical processes.
4. Thus, science can model what pain feels like.

It is common to suppose that knowing a scientific model of pain does not give knowledge of what pain feels like. In contrast, according to Naturalistic Monism, knowing the pain-model *does* give knowledge of what pain feels like, but only knowledge in terms of the scientific model, which is distinct from concrete pain. What concrete pain feels like is simply the happening of pain in us, a process that science aims to model with the pain-model. We can say that pain is nothing but “T-interaction” (in a *de re* sense, meaning that “T-interaction” *refers* to pain), but this does not amount to reducing concrete pain into the theoretical T-interaction-model. Instead, pain, including what we call its “what-it-is-likeness”, is the concrete phenomenon that science *models as* T-interaction. The take-home message is that we need not postulate the existence of any properties (“qualia”) for the what-it-is-likeness of pain over and above the properties described by the scientific pain-model. Once we have a scientific model of pain that is perfectly isomorphic with pain as experienced, we have a scientific model of pain’s phenomenology. For instance, the experienced intensity or sharpness of pain could be modelled as parameters in the pain-model that correspond to such qualities—any change in the experienced sharpness of pain corresponds to a change in the corresponding parameter (see section 4.1.). However, this does not mean that pain itself would be just the theoretical parameters; it is what the parameters model.^3^

## 4. Studying consciousness scientifically

We have argued, first, that consciousness should be viewed as a concrete physical process in the world in the same sense as lightning, photosynthesis, and life—experiences are of the same ontological type as all physical phenomena, there is only difference in complexity. Second, we have stressed that scientific models are themselves mental phenomena and always distinct from the phenomena they model. Because experiences are concrete phenomena, they can causally interact with other physical phenomena, including observers and their measuring devices. Science can model experiences as CMCs. For a neuroscientist, the experience of a subject is the “external” phenomenon that affects the scientist’s measuring devices, and which they aim to describe with the CMC-model. From the subject’s perspective, their experience is an “internal” phenomenon, part of their subjective realm.

We will next illustrate Naturalistic Monism with the help of a neuroscientific example. Consider a visual experience of a cat (Figure 1). On our approach, a subjective cat experience is the physical process in the world that underlies neuroscientific observations that we interpret in terms of the CMC_cat_-model (a model of the constitutive mechanisms of a cat experience). Thus, the terms “cat experience” and “CMC_cat_” refer to the same concrete phenomenon.^4^ Whereas the term “experience of cat” is typically used to refer to an experience when it is part of a subject’s consciousness, the neuroscientific term “CMC_cat_” is used in scientific contexts, where it refers to the experience as the external phenomenon that causes scientific observations. Experiences are complex physical processes that are composed of lower-level phenomena, and the composition can be described scientifically. Science can describe how experiences are composed of lower level processes in the same way as it explains how hydrogen and oxygen form water. This framework is summarized in Figure 1.

**Figure 1.**
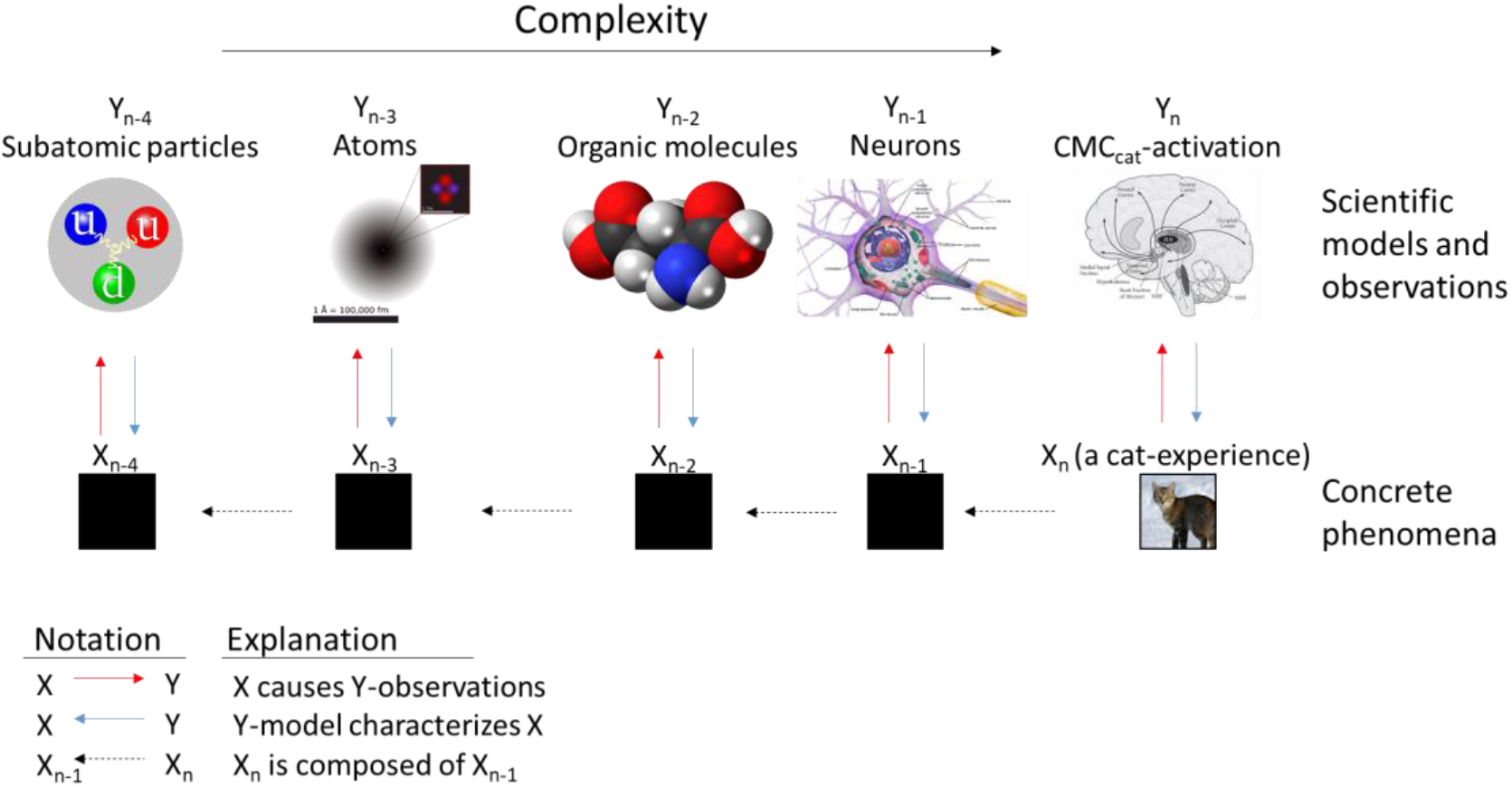
A schematic presentation of Naturalistic Monism. Empirical observations (upper row) are caused by concrete physical phenomena (red arrows), and empirical models characterize the concrete physical phenomena (blue arrows). Complex physical phenomena are composed of more basic level phenomena (dashed arrows). Experiences, as depicted by the picture of a cat, are concrete physical phenomena, and hence can be empirically observed and described by scientific models. Because experiences are complex physical phenomena, they are composed of lower-level constituent processes. The lower-level constituents of experiences are shown in black because they are not part of our consciousness; they can only be characterized empirically. (All images under Creative Commons license.)

The cat-experience is depicted in Figure 1 as a complex physical process that is composed of lower-level constituent processes. This can be empirically modelled by defining how the CMC_cat_-process is related to its constituent processes. It is often argued that consciousness cannot be fully explained by analyzing the interactions between its lower-level constituents (this is the so-called “combination problem” that RM faces; see Chalmers, 2017). This argument conflates phenomena with their scientific models: one can never “reduce” any phenomenon to its description or descriptions of its lower level processes (reduce X_n_ to Y_n_ or Y_n-1_ in Figure 1); one can only reduce descriptions to other descriptions (reduce Y_n_ to Y_n-1_). To give an example, one can *explain* how the functioning of an amoeba is based on the interactions between the molecules that constitute it, and in this sense reduce the *scientific model* of the living amoeba to *models* of lower-level phenomena, such as interactions between proteins, enzymes, and so on. This does not mean that the actual, concrete amoeba is “reduced” to anything else; the concrete amoeba *is* the complex system composed of the interaction between molecules, enzymes, and so on. In the same way, one can *model* how an experience is based on lower level processes (how Y_n_ is based on Y_n-1_), but this does not amount to “reducing” the concrete experience (X_n_) into a description. When we explicate how a cat-experience is composed of lower level processes (the lower axis in Figure 1), we do it in terms of describing how the CMC_cat_ is composed of lower-level processes, such as the activation of individual neurons (the upper axis in Figure 1). Crucially, when we scientifically reduce an experience into lower-level processes, we do so only descriptively. “Reducing” the concrete experience—literally tearing it to parts—would amount to destroying it. The crucial point here is that if we accept that scientific models of experiences are literally models of experiences, then it is possible to scientifically model how subjective experiences are based on lower-level processes.

The black boxes in Figure 1 might appear to motivate some kind of mysticism about the nature of (non-mental) physical phenomena, similar to Strawson’s “intrinsic nature” that cannot be empirically captured. However, this mysticism is inherent to all science: we can never know the nature of the world as it is in itself, we can only model it based on how it affects our senses. In Figure 1, the lower level boxes are mostly black to emphasize that the phenomena exist independently and outside of how we scientifically model them, but this does not imply that they are *completely* epistemologically opaque to us: we *can* know their nature, but *only* through observations and models. Moreover, it is important to notice that whether a box is black or not depends on perspective: my consciousness is not a black box for me, but it is for you, because your knowledge of my consciousness is based solely on observations and models (your model of my mind).

### 4.1. Towards an empirical model of consciousness

Because Naturalistic Monism implies that an experience is a concrete physical phenomenon, and that all the properties of experiences are physical properties, it must be possible to observe and scientifically model all the characteristics of an experience. Hence, based on the current framework, empirical science can study consciousness, not just its correlates. This is an important upshot, as it collapses the problem of consciousness into a standard problem of science.

Specifically, we suggest that scientific models of consciousness should describe the following^5^ (Figure 2):

1. *Constituents*. Because consciousness is a complex physical process, science should describe its constituents. This explanation describes what consciousness is “made of”.
2. *Isomorphy*. A complete model of consciousness should accurately describe all the details about the contents of consciousness, including its phenomenology.
3. *Etiology*. Understanding the etiological basis of consciousness, from immediately preceding causes to its phylogenetic basis, is crucial for understanding phenomenology.
4. *Causal power*. Being a concrete physical phenomenon, consciousness can causally interact with other physical processes inside the brain. A key part of explaining consciousness is explaining what it can do.

**Figure 2.**
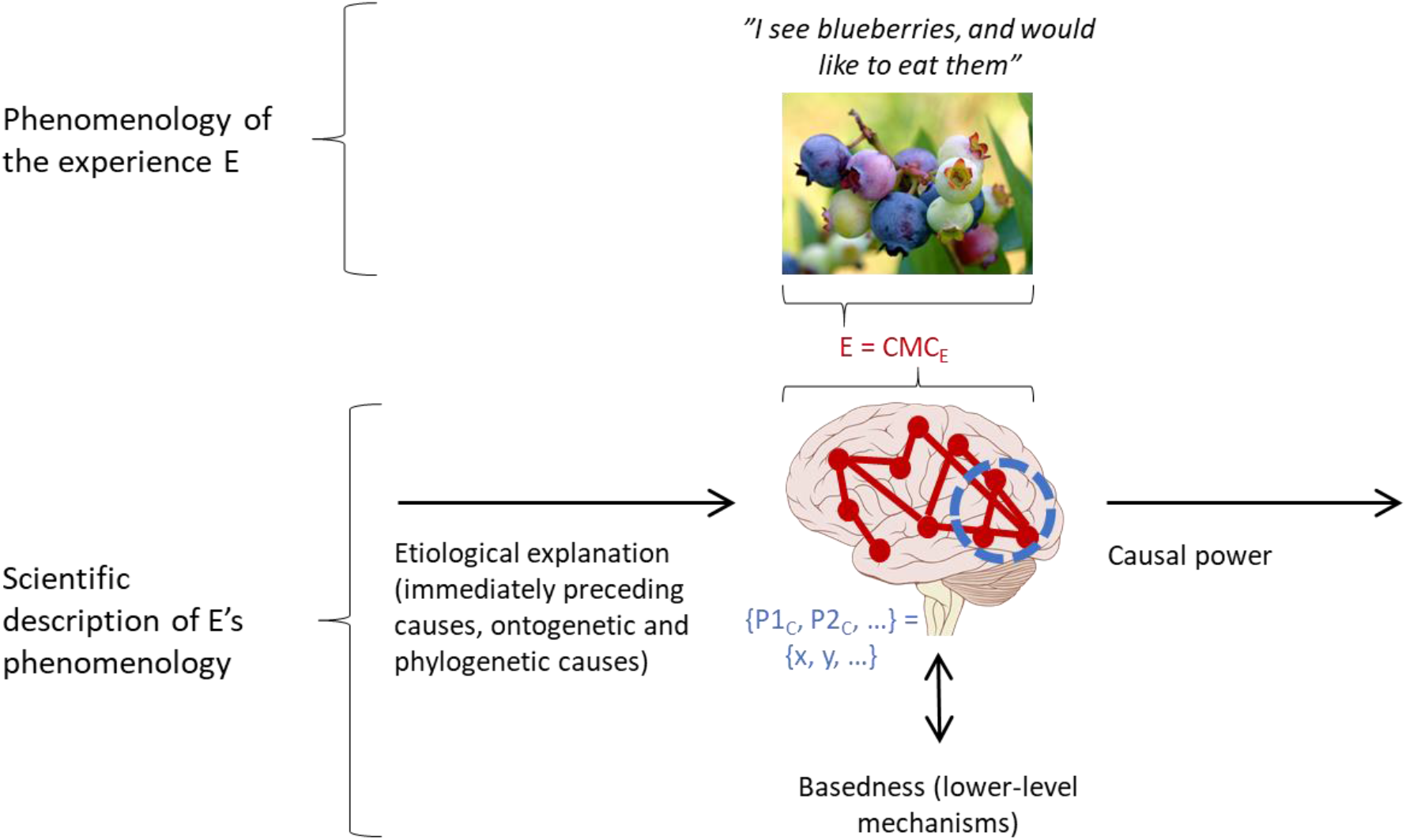
Understanding phenomenology empirically. Phenomenology of experience E is depicted in the figure by the picture of blueberries. We assume that at this moment the subject experiences seeing blueberries, and feels an urge to eat them. The scientific model of E is here depicted by the image of the brain (CMC_E_), where color is represented as a set of color parameters {P1_C_, P2_C_, …} that can have different values {x, y, …}. Our key claim is that the CMC_E_ is a model of the experience E. Hence, CMC_E_ and E, although appearing to be different, are isomorphic, and E can be described by a detailed empirical model (CMC_E_). To understand the “mechanism of consciousness”, science must describe the lower-level processes that constitute it, and describe the preceding causes of E (etiology; this include e.g. phylogeny). Causal power here refers to the neural processes and behavior that the experience E (empirically described as CMC_E_) causes. [All images under Creative Commons license.]

#### 4.1.1. Lower-level constituents

According to Naturalistic Monism, a successful empirical theory of consciousness must specify the constituent processes that are necessary for consciousness, but which are not themselves conscious until they interact in some specific way to form the CMC. The constituents are for CMC what hydrogen and oxygen are for water: hydrogen and oxygen do not cause water, they constitute it. The constituents of consciousness are often called the “mechanism of consciousness”: Which neural processes determine whether an individual is conscious or unconscious? Which neural processes determine whether a visual stimulus will be consciously perceived? Examples of theoretical attempts to answer such question include the global workspace model (GWT; Baars, 1988; Dehaene & Changeux, 2011), integrated information theory of consciousness (IIT; Oizumi, Albantakis, & Tononi, 2014), and higher-order thought theory of consciousness (Lau & Rosenthal, 2011). These theories aim to describe how consciousness “emerges” from neural activity. According to Naturalistic Monism, these theories aim to describe the CMC. Crucially, because the CMC is a model of consciousness, explaining the constituents of CMC amounts to explaining the constituents of consciousness itself—not just something that correlates with consciousness.

Consider the GWT, for example. According to the theory, consciousness is a process where a large number of individual modules share their information with each other to enable flexible behavior and maintenance of information (Baars, 1988). According to the neural model of the theory, this corresponds to synchronous interactions between specific neural populations in frontal, parietal and sensory cortical areas (Dehaene & Changeux, 2011). These individual processes are not conscious alone, but become conscious when they interact in the way that the theory describes. These constituent processes are further based on lower-level processes: oscillatory activity requires a certain balance of excitatory and inhibitory neurons, neural activity requires a resting membrane potential, neurons are made up of specific molecules, and so on. The constituents can, in principle, be traced to fundamental physical entities; in this sense, the CMC (the experience) is continuous with other physical phenomena.

GWT does not make claims about how the subjective aspects of consciousness (phenomenology) are enabled by activation of the global workspace, but according to Naturalistic Monism, the concrete phenomenon whose functional aspects GWT models *is* consciousness and has a certain phenomenology for the subject in whom it happens. Accordingly, the functional properties modelled by GWT are not independent of phenomenal consciousness; they are the functional aspects *of* phenomenal processes (causal power of consciousness is discussed further in section 4.1.4.).

#### 4.1.2. Isomorphism

Because the phenomenology of an experience is a concrete physical process, any change in phenomenology can in principle be observed and modelled by a specific change in the model. Thus, a complete CMC_E_-model of an experience E must be perfectly isomorphic with the experience: there must be a one-to-one correspondence between the empirically observed physical phenomenon (CMC_E_) and the subjective experience E (Fingelkurts & Fingelkurts, 2004; Revonsuo, 2000). This means that an empirical theory of consciousness, if sufficiently detailed, can provide a complete scientific description of an experience and its phenomenology. To do this, the theory must be able to define which empirically observed physical processes correspond to different contents of experiences, and how these contents combine to form complex experiences. For instance, from the perspective of conscious vision, the requirement of isomorphy means that the model must predict the phenomenology of vision from the egocentric perspective, not from the retinotopically-centered frame of reference (Land, 2012). According to the integrated information theory of consciousness (Oizumi et al., 2014) the structure of recurrent networks (“qualia space”, to use IIT’s terminology) defines phenomenology. According to Naturalistic Monism, such models are descriptions of the concrete physical phenomenon that is consciousness.

The reader may, at this point, insist that isomorphic descriptions of CMCs would not explain the “qualitative feelings” of experiences, or answer why certain neuronal activity feel like something, and hence the description would not be an explanation of phenomenology. Such counter-arguments are ill-posed, because they conflate concrete phenomena with their scientific models. For instance, what blue feels like for S is the happening of CMC_blue_ in S, a concrete process that can be perfectly modelled scientifically—but just *modelled*. Grasping the CMC_blue_-model affords knowledge of how an experience of blue can be scientifically *described*, whereas what blue feels like is just the happening of the concrete phenomenon, which is distinct from the scientific model. No description alone can ever cause the happening of the concrete phenomenon it describes, but this does not imply that it could not perfectly describe the phenomenon and its happening in a subject. Science can model the constituents, etiology, etc. of blue experiences, and there is no further question of why blue (or the corresponding neural process) feels like blue. Feeling like blue is part of the nature of blue experiences, determined by their constituent processes, just like having a certain molar mass is part of the nature of helium. Science can model what blue feels like and how that feel is determined by lower-level processes, just like it can model the molar mass of helium and its constituent processes.

#### 4.1.3. Etiology

In trying to explain what must happen in the brain so that, for instance, visual information crosses the threshold to consciousness, neuroscience examines the causal chain of events that lead to it (Dehaene & Changeux, 2011; Railo, Koivisto, & Revonsuo, 2011). In other words, they describe the etiology of conscious vision. The aim is to disentangle the CMC from other processes that correlate with conscious perception, but do not constitute it (e.g. Aru, Bachmann, Singer, & Melloni, 2012). For example, neural processes that precede the presentation of a stimulus help to predict whether or not the stimulus is consciously perceived – in other words, prestimulus activity correlates with the subsequent conscious perception (e.g. Iemi, Chaumon, Crouzet, & Busch, 2016). However, these prestimulus correlates of conscious vision are obviously “just correlates”, and the neural processes that constitute the conscious perception of the stimulus happen later. One of the challenges of empirical science is to characterize this etiological chain of events that leads to conscious vision, that is, the formation of the CMC that corresponds to experience of that stimulus.

In addition to explaining the immediately preceding causes that lead to consciousness, science can describe longer time scales such as individual development, or evolution (see Revonsuo, 2006). A complete theory of consciousness should explain how consciousness has evolved from earlier products of evolution. For example, the phenomenology of visual perception could be accounted wholly in empirical terms by characterizing how frequently different stimulus combinations have been encountered during the evolution (Purves, Wojtach, & Lotto, 2011). Similarly, pain can be considered as a complex biological adaptation that has helped organisms to avoid potentially damaging stimuli, and whose bio-chemical basis could possibly be traced back to single celled organisms (Bray, 2009). According to Naturalistic Monism, this does not mean that only the functional aspects of pain (or visual perception) would be products of evolution, but also that the different types of subjective unpleasant sensations we call “pain” (or visual phenomenology) are products of evolution. This view is built on the assumption that consciousness is not an epiphenomenon (cf. Jackson, 1982), but instead that it has power to influence other physical processes.

#### 4.1.4. Causal power

The idea that experiences are something different from other physical phenomena is deep-rooted in philosophy and science. For instance, Edelman (2003), who builds an elegant theory of consciousness on a bio-physical foundation, argues that conscious states are “informational even if not causal” (p. 5523). This way, consciousness easily appears epiphenomenal, which is strongly at odds with our subjective experience as well as the empirical scientific assumption that human consciousness is a product of evolution, and plays a causal role in behavior. Naturalistic Monism ties together the phenomenal and functional aspects of consciousness. Because consciousness is a physical process, it can causally interact with other phenomena in the brain and drive the organism’s behavior. We propose that Naturalistic Monism may help to dismantle some stubborn conflicts about whether consciousness should primarily be described in functional or phenomenal terms (Block, 1995; Carter et al., 2018; Dehaene & Changeux, 2011). While it is an empirical question to find out whether specific cognitive functions (e.g. global access or error detection; Dehaene, Lau, & Kouider, 2017) are necessary for consciousness, our framework implies that experiences never take place in a causal void. An experience E (CMC_E_) always causally interacts with other processes in the brain, both conscious and non-conscious: it can cause, for instance, thoughts, emotions, bodily states, or actions, and interact with the outside world. One of the goals of empirical science of consciousness is to explain what gives conscious processes their power to potentially elicit fast and accurate actions, and flexible voluntary behavior^6^. Theories such as the Global Workspace theory are explicitly built around this foundation (Dehaene & Changeux, 2011). According to Naturalistic Monism, such “functional explanations” can provide key insights into the phenomenal qualities of experiences as well (cf. the example about pain, in previous subsection).

## 5. Philosophical implications

In this section, our aim is to briefly address the main philosophical implications of Naturalistic Monism. The discussion is not intended as exhaustive, but rather as pointing towards future research.

### 5.1. Relativity of “subjective” and “objective”

Naturalistic Monism implies that terms like “subjective”/”objective”, or “first-person”/”third-person” are relative. When a subject S1 perceives an apple, the apple is objective and third-person observable. The experience of the apple in S1 is subjective or first-person observable for S1, but because it is a natural phenomenon, it is third-person observable for a neuroscientist (S2). The neuroscientist’s observations and theoretical models of the neural mechanisms of S1’s apple-experience, in turn, are phenomena in the scientist’s mind/brain and (at least partly^7^) subjective or first-person observable for the neuroscientist—the scientist is *aware of* grasping the model. The scientist’s model is in principle objectively or third-person observable for a “metaneuroscientist” (S3), and so on *ad infinitum*. In short: any subject’s experience is another subject’s external phenomenon.

Husserl’s notions of phenomenological and natural attitude can be used to illustrate this relativity. In everyday life, we typically endorse the natural attitude, focusing on external objects instead of our experiences of them. When I perceive an apple, my consciousness is directed at the apple and I am not necessarily aware of the details of my experience *per se*, such as what the greenness of the apple feels like. In Hurssel’s terminology this is called the natural attitude. In contrast, when I endorse the phenomenological attitude, I focus on my apple-experience *as an experience*. A scientist, in turn, is concerned with the external phenomena they study and thus automatically endorse the natural instead of the phenomenological attitude—they rarely focus on the nature of their scientific models as conscious phenomena in their minds. However, even the conceptual aspects of a scientific model—e.g. that the helium atom has two protons—are conscious phenomena in the mind of the scientist when they grasp the model. As such, the scientist can endorse the phenomenological attitude towards them and consider them as experiences. This illustrates that we can never escape our own point of view: even a scientific model is only a representation in the scientist’s mind. However, even though a scientist S1 cannot themselves “step outside” their model, an external observer S2 can: in principle, such a “meta-observer” could investigate the relationship between an object X and the scientist S1’s mental model of it from an external perspective, treating both the object the model as external phenomena (“noumena” in Husserl’s terms). But the meta-observer S2 would then be confined to their own perspective (and so on for any further meta-observer).

### 5.2. Philosophies of science and consciousness

If we consider consciousness as a concrete physical phenomenon, then the question about the relationship between an experience and its mature scientific model comes at least partly down to which philosophy of science we endorse (in addition to the details of the model itself). A scientific model is always distinct from the concrete phenomenon it models, and there is always a gap between the two in this sense. How wide we take the gap to be depends on whether we endorse realism or antirealism about science. Roughly, according to realism, mature scientific theories can be true, and truthfully describe the nature of the world (Psillos, 1999). Thus, the epistemic gap between a phenomenon and its mature scientific model is non-existent or narrow.^8^ According to antirealism, on the other hand, we always have to remain agnostic about the truth of scientific models. For instance, according to constructive empiricism (van Fraassen, 1980), scientific theories can be considered as *empirically adequate* if they are isomorphic with observations, but we can never say whether they are literally true. We could say that, on this view, there is always an indefinitely wide epistemic gap between a phenomenon and its model, or that it makes no sense to speak of such a gap.

Scientific realism implies that if we said that our experiences involve purely qualitative aspects that cannot be modelled structural-relationally, we would be mistaken. According to scientific realism, *all* natural phenomena can be truthfully modelled structural-relationally. On this view, what we call a “qualitative feeling” can be truthfully modelled as a structural-relational process. Importantly, this approach does not deny that experiences feel like something; it only holds that the feel can be *modeled* purely structural-relationally. If we *characterized* the feel as something purely qualitative, we would be mistaken—not about the feel itself, but instead about how to describe it. Importantly, terms like “qualitative” or “feeling” are only concepts which refer to certain concrete processes that happen in us. The processes are the way they are, no matter how we refer to them. Scientific realists would hold that experiences can be truthfully modelled completely in structural-relational terms, but this does not, and cannot, amount to eliminating any aspects of the experiences themselves. It may be difficult to see how, say, the redness of red could be a structural-relational process, given that introspection about redness does not reveal that it has such nature. However, from the fact that a model M applies to an experience E, it does not follow that a subject experiencing E would introspectively see that M applies to it (cf. Dennet, 1991).

If we endorse antirealism, such as van Fraassen’s empiricism, we can only say whether a neuroscientific theory of consciousness is compatible with observations, not whether it “captures the nature” of what it models. It could be said that, for the antirealist, there is *always* an indefinitely wide epistemic gap between *any* concrete phenomenon and its scientific model, and that experiences are not different in this respect. It is noteworthy that Naturalistic Monism is not committed to either realism or antirealism; this is an independent question. Naturalistic Monism minimally implies that there is always *distinctness* between a phenomenon and its model, but whether we take there to be an *epistemic* gap may depend on which philosophy of science we endorse.

Detailed discussion of how consciousness is related to different philosophies of science is beyond the scope of this paper. The main point is that if we consider experiences as being of the same (ontological) type as any other physical phenomenon, then the question about the relationship between an experience and its scientific model boils down to the general question about the relationship between *any* phenomenon and its model.

### 5.3. Panpsychism

According to Naturalistic Monism, experiences are concrete physical phenomena that are based on lower-level physical processes, and thereby ontologically continuous with them. Does it follow that *all* physical phenomena are of an “experiential nature”, because they are of the same ontological type as experiences (cf. the black boxes in Figure 1)? In our view, this is not a necessary conclusion. An experience E can be scientifically modelled as CMC_E_, a system-level phenomenon in the brain that is based on the activity of individual neurons. In this sense, consciousness is continuous with other physical phenomena, all the way down to subatomic particles. However, insofar as individual neurons (not to speak of quarks) alone do not realize the CMC_E_-process, they are not conscious, at least in the same sense as we are. On this line of argumentation, it is possible for consciousness to emerge from something that is not conscious, and it is the task of empirical theories of consciousness to describe exactly how this happens (but see Strawson, 2008b).

On the other hand, we have no non-empirical or non-model-mediated access to the nature of other physical phenomena besides our experiences. We can extrapolate from our own case and infer that at least cats and cows are likely to have similar experiences as us, given that they show similar behaviors and have similar neuroanatomy. It is more difficult to say anything about the possible inner lives of insects, amoebas, or quarks. This difficulty, however, stems from the limits of science, not from consciousness being a special type of phenomenon.

### 5.4. Zombies, Mary, and the “hard problem”

Philosophical zombies (Chalmers, 1996) are creatures that appear just like normal human beings in every functional and observable respect, but for whom “the lights are out”: it does not feel like anything to be a zombie. When we imagine zombie-pain, we imagine something that would cause pain-observations and fit our scientific model of pain, but which would not be real pain. If such fake-pains are conceivable, then arguably we could conceive of any fake natural phenomena. For instance, we can imagine that we live in a massive computer simulation, where our observations of electrons are not caused by real electrons, but instead by computer algorithms. If such imaginings are possible at all, what they teach us is that our scientific models and observations are always distinct from the concrete phenomena that underlie them. It does not tell us anything about the nature of experiences or electrons, except for that they are concrete phenomena and independent of their scientific models and observations.

The same reasoning can be applied to the Mary case, a neuroscientist who has never experienced colors (Jackson, 1986). Mary knows everything about the visual color system in the brain, but she does not know what it is like to see red. The conclusion that the thought experiment is supposed to demonstrate is that there are some facts about experiences that cannot be described by scientific models. But the conclusion is false: Mary *does* know what it is like to see red, but only in terms of a scientific model. What it is like for S to see red is the happening of a red experience in S. Because science can model such processes, it can thereby model what it is like to see red. When Mary sees red for the first time, the concrete phenomenon whose model she already knew *happens* in her. What the Mary case teaches us is that a concrete phenomenon can be quite different from what a scientific model says it is. But this does not imply that there would be some properties (“qualia”) that cannot be scientifically modelled, it only shows that a concrete phenomenon is distinct from its model.

## 6. Conclusions

We have argued that once we clearly distinguish between concrete phenomena and their scientific models, the epistemic gap – why our experiences appear so different from neural activity – dissolves into distinctness between the model and the phenomenon. From the perspective of Naturalistic Monism, science can observe and model consciousness, not just its correlates. We hope that our proposal helps to establish a solid ground for the empirical science of consciousness, as it collapses the problem of consciousness into a standard problem of science. Scientific explanation never affords us knowledge of the nature of phenomena in themselves. However, if we accept that our subjective experiences are concrete physical phenomena, then science can provide a complete and exhaustive *model* of experiences, including what they “feel like”.

## 7. Acknowledgements

J.J. was supported by the Abo Akademi University Endowment (the BrainTrain project and a personal grant) and H.R. by the Academy of Finland (grant #308533). We thank Antti Revonsuo, Valdas Noreika, and Adam Bricker for helpful comments and discussions.

## Footnotes

1 The phenomenal concepts strategy (PCS) aims to do just this: account for the epistemic gap by analyzing the referential mechanisms of phenomenal and scientific concepts (see above). Our approach can be considered as largely compatible with PCS, but we view the question from a different angle: we focus not on semantics, but instead on the nature of science and the relationship between scientific models and the concrete phenomena they model.

2 This is a simplification, because conceptualizing an experience can modify it—for instance, verbalizing an emotion can weaken its intensity (e.g. Torre & Lieberman, 2018).

3 Dennet (1991) makes a similar case, but he does not clearly distinguish between concrete phenomena and their theoretical models. Arguably, this has led people to accuse him of being an eliminativist about experiences. However, it is one thing to be an eliminativist about “qualia”, a theoretical concept which holds that experiences involve completely non-structural, non-relational, and non-dispositional properties, and a completely different thing to be an eliminativist about concrete experiences. Dennett is an eliminativist about the former, but not about the latter. On our view, he would simply claim that experiences can be *modelled* completely functionalistically, but this does not amount to claiming that the modelled phenomena themselves would not feel like something (in the everyday sense of “feeling like something”).

4 Here it is important to distinguish between experience- and CMC-*types*, and their *tokens*, such as a particular cat-experience. We do not wish to take a stand on this question here, but we assume that at least each experience token is identical with what its complete neuroscientific model token refers to. Types are more complex because they are always abstractions; we could question whether there can exist any cat-experience type. Models, in turn, are themselves abstractions based on observations of tokens, and cannot be considered as strictly identical with any particular experiences. For a discussion on the type-token distinction in physicalism, see e.g. Stoljar (2001).

5 It may be impossible for any *single* theory of consciousness to fulfill all the listed desiderate; for instance, we might need one theory about the functions of consciousness and another (compatible) theory about isomorphy. This is similar to what Hawking and Mlodinow (2010) suggest with respect to the possibility of a Grand Unified Theory in physics: it might be impossible to model the world in terms of a single theory, but instead we might need a set of mutually compatible theories.

6 While stimuli that are not consciously experienced may influence individual’s behavior (Cowey, 2010; Hannula, Simons, & Cohen, 2005), it is important to note that the causal power of unconscious information on behavior is very limited compared to conscious perception (see, Revonsuo, Johanson, Wedlund, Chaplin, 2000, for discussion about the empirical possibility, and potential empirical examples of zombie behavior). Moreover, recent research has also shown that early studies overestimated the power of unconscious processes to influence behavior (Ffytche & Zeki, 2011; Newell & Shanks, 2014; Overgaard, 2011; Peters & Lau, 2015)

7 Any experience involves also subconscious processes, and highly complex conceptual mental representations such as scientific models may involve many subconscious processes.

8 Naturalistic Monism implies that the model and the modeled phenomenon are always *distinct*, even if we would endorse realism and say that the model is “true”. On this approach, between a model and a phenomenon there would be a gap in the sense of distinctness but not necessarily an *epistemic* gap. Moreover, even though Naturalistic Monism implies that we cannot know the modeled phenomenon as it is in itself, our descriptions of it can still be considered as “true” if, for example, the components of the model are structurally isormophic with properties of the modelled phenomenon (cf. Worrall, 1989).

## References

Alter, T., & Nagasawa. (2012). What is Russellian Monism. Journal of Consciousness Studies, 19(9–10).

Aru, J., Bachmann, T., Singer, W., & Melloni, L. (2012). Distilling the neural correlates of consciousness. Neuroscience and Biobehavioral Reviews. https://doi.org/10.1016/j.neubiorev.2011.12.003

Baars BJ, & Baars, B. J. B. (1988). A cognitive theory of consciousness. A Cognitive Theory of Consciousness. https://doi.org/10.1108/09513551111163639

Balog, K. (2012). In Defense of the Phenomenal Concept Strategy1. Philosophy and Phenomenological Research, 84(1), 1–23. https://doi.org/10.1111/j.1933-1592.2011.00541.x

Block, N. (1995). On a confusion about a function of consciousness. Behavioral and Brain Sciences. https://doi.org/10.1017/S0140525X00038188

Bray, D. (2009). Wetware: A Computer in Every Living Cell. Yale: Yale UP.

Carter, O., Hohwy, J., Van Boxtel, J., Lamme, V., Block, N., Koch, C., & Tsuchiya, N. (2018). Conscious machines: Defining questions. Science. https://doi.org/10.1126/science.aar4163

Chalmers, D. (1996). The Conscious Mind: in Search of a Fundamental Theory. Oxford: Oxford UP.

Chalmers, D. (2016). Panpsychism and panprotopsychism. In G. Büntrup & L. Jaskolla (Eds.), Panpsychism: Contemporary Perspectives. Oxford: Oxford UP.

Chalmers, D. (2017). The combination problem for panpsychism. In Panpsychism. Contemporary Perspectives. Oxford: Oxford UP.

Cowey, A. (2010). The blindsight saga. Experimental Brain Research. https://doi.org/10.1007/s00221-009-1914-2

Dehaene, S. (2014). Consciousness and the Brain - Deciphering How the Brain Codes Our Thought. New York: Penguin Books.

Dehaene, S., & Changeux, J. P. (2011). Experimental and Theoretical Approaches to Conscious Processing. Neuron. https://doi.org/10.1016/j.neuron.2011.03.018

Dehaene, S., Lau, H., & Kouider, S. (2017). What is consciousness, and could machines have it? Science. https://doi.org/10.1126/science.aan8871

Dennet, D.. (1991). Consciousness Explained. Boston: Little, Brown and Co.

Dennett, D. C. (2018). Facing up to the hard question of consciousness. Philosophical Transactions of the Royal Society B: Biological Sciences, 373(1755), 20170342. https://doi.org/10.1098/rstb.2017.0342

Eddington, A. (1929). The Nature of the Physical World. Cambridge: Cambridge UP.

Edelman, G. M. (2003). Naturalizing consciousness: A theoretical framework. Proceedings of the National Academy of Sciences. https://doi.org/10.1073/pnas.0931349100

Edelman, G. M., & Tononi, G. (2000). A Universe of Consciousness: How Matter Becomes Imagination. Basic Books.

Ellis, B. (2001). Scientific Essentialism. Cambridge: Cambridge UP.

Ffytche, D. H., & Zeki, S. (2011). The primary visual cortex, and feedback to it, are not necessary for conscious vision. Brain. https://doi.org/10.1093/brain/awq305

Fingelkurts, A. A., & Fingelkurts, A. A. (2004). Making complexity simpler: Multivariability and metastability in the brain. International Journal of Neuroscience. https://doi.org/10.1080/00207450490450046

Francis Crick and Chri.dof Koch. (1990). Towards a neurobiological theory of consciousness. The Neurosciences. https://doi.org/10.1109/INEC.2010.5424508

Friston, K. (2010, February 13). The free-energy principle: A unified brain theory? Nature Reviews Neuroscience. Nature Publishing Group. https://doi.org/10.1038/nrn2787

Goff, P. (2017). Consciousness and Fundamental Reality. Oxford: Oxford UP.

Hannula, D. E., Simons, D. J., & Cohen, N. J. (2005). Imaging implicit perception: promise and pitfalls. Nature Reviews Neuroscience. https://doi.org/10.1038/nrn1630

Hawking, S. (1988). A Brief History of Time. New York: Bantam Books.

Hawking, S., & Mlodinow, L. (2010). The Grand Design. New York: Bantam Books.

Hiddleston, E. (2019). Dispositional and categorical properties, and Russellian Monism. Philosophical Studies, 176(1), 65–92. https://doi.org/10.1007/s11098-017-1006-2

Hobson, J. A., & Friston, K. J. (2014). Consciousness, Dreams, and Inference: The Cartesian Theatre Revisited. Journal of Consciousness Studies, 21(1–2), 6–32.

Howell, R. (2015). The Russellian Monist’s Problems with Mental Causation. The Philosophical Quarterly, 65(258), 22–39. https://doi.org/10.1093/pq/pqu058

Iemi, L., Chaumon, M., Crouzet, S. M., & Busch, N. A. (2016). Spontaneous Neural Oscillations Bias Perception by Modulating Baseline Excitability. The Journal of Neuroscience. https://doi.org/10.1523/jneurosci.1432-16.2016

Ismael, J. (1999). Science and the phenomenal. Philosophy of Science, 66(3), 351–369.

Jackson, F. (1982). Epiphenomenal qualia. The Philosophical Quarterly, 32(127), 127–136.

Jackson, F. (1986). What Mary Didn’t Know. The Journal of Philosophy, 83(5), 291–295. https://doi.org/10.2307/2026143

Kind, A. (2015). Pessimism about Russellian monism. In T. Alter & Y. Nagasawa (Eds.), Consciousness in the Physical World: Perspectives on Russellian Monism (pp. 401–421). Oxford: Oxford UP.

Koch, C., Massimini, M., Boly, M., & Tononi, G. (2016). Neural correlates of consciousness: Progress and problems. Nature Reviews Neuroscience. https://doi.org/10.1038/nrn.2016.22

Land, M. F. (2012). The operation of the visual system in relation to action. Current Biology. https://doi.org/10.1016/j.cub.2012.06.049

Lau, H., & Rosenthal, D. (2011). Empirical support for higher-order theories of conscious awareness. Trends in Cognitive Sciences. https://doi.org/10.1016/j.tics.2011.05.009

Laureys, S. (2005). The neural correlate of (un)awareness: Lessons from the vegetative state. Trends in Cognitive Sciences. https://doi.org/10.1016/j.tics.2005.10.010

Loar, B. (1990). Phenomenal states. Philosophical Perspectives, 4, 81–108.

Michel, M., Fleming, S. M., Lau, H., Lee, A. L. F., Martinez-Conde, S., Passingham, R. E., … Liu, K. (2018). An informal internet survey on the current state of consciousness science. Frontiers in Psychology. https://doi.org/10.3389/fpsyg.2018.02134

Montero, B. (2015). Russellian physicalism. In T. Alter & Y. Nagasawa (Eds.), Consciousness in the Physical World (pp. 209–223). Oxford: Oxford UP.

Nagel, T. (1979). What it is like to be a bat? In Mortal Questions (pp. 165–180). Cambridge: Cambridge UP.

Newell, B. R., & Shanks, D. R. (2014). Unconscious influences on decision making: A critical review. Behavioral and Brain Sciences. https://doi.org/10.1017/S0140525X12003214

Oizumi, M., Albantakis, L., & Tononi, G. (2014). From the phenomenology to the mechanisms of consciousness: integrated information theory 3.0. PLoS Computational Biology, 10(5). https://doi.org/10.1371/journal.pcbi.1003588

Overgaard, M. (2011). Visual experience and blindsight: A methodological review. Experimental Brain Research. https://doi.org/10.1007/s00221-011-2578-2

Papineau, D. (2002). Thinking about Consciousness. Oxford: Oxford UP.

Peters, M. A. K., & Lau, H. (2015). Human observers have optimal introspective access to perceptual processes even for visually masked stimuli. ELife. https://doi.org/10.7554/eLife.09651

Physical. In OxfordDictionaries.com. Retrieved from https://en.oxforddictionaries.com/definition/physical

Psillos, S. (1999). Scientific Realism: How science tracks truth. London: Routledge.

Purves, D., Wojtach, W. T., & Lotto, R. B. (2011). Understanding vision in wholly empirical terms. Proceedings of the National Academy of Sciences. https://doi.org/10.1073/pnas.1012178108

Railo, H., Koivisto, M., & Revonsuo, A. (2011). Tracking the processes behind conscious perception: A review of event-related potential correlates of visual consciousness. Consciousness and Cognition. https://doi.org/10.1016/j.concog.2011.03.019

Revonsuo, A.; Johanson, M.; Wedlund, J-E.; Chaplin, J. (2000). The zombies among us. Consciousness and automatic behavior. In A. Rossetti, Y.; Revonsuo (Ed.), Beyond Dissociation. Interaction between dissociated implicit and explicit processing. (pp. 331–3351). Amsterdam: John Benjamins.

Revonsuo, A. (2000). Prospects for a Scientific Research Program on Consciousness. In T. Metzinger (Ed.), Neural Correlates of Consciousness. Cambridge, MA: MIT Press.

Revonsuo, A. (2006). Inner Presence: Consciousness as a Biological Phenomenon. Massachusetts: MIT Press.

Russell, B. (1927). The Analysis of Matter. London: Kegan Paul.

Schneider, S. (2017). What breathes fire into the equations? Journal of Consciousness Studies, 24(9–10).

Seager, W. (2006). The “intrinsic nature” argument for panpsychism. Journal of Consciousness Studies, 13(10–11), 129–145.

Searle, J. R. (2002). Why I am not a property dualist. Journal of Consciousness Studies, 9(12), 57–64. Retrieved from https://philpapers.org/rec/SEAWIA

Stoljar, D. (2001). Physicalism. In Stanford Encyclopedia of Philosophy. Retrieved from https://plato.stanford.edu/entries/physicalism/

Strawson, G. (2008a). Real materialism. In Real Materialism and Other Essays (pp. 19–52). New York: Oxford UP.

Strawson, G. (2008b). Real Materialism and Other Essays. Oxford: Oxford University Press.

Strawson, G. (2008c). Realistic monism: Why physicalism entails panpsychism. In Real Materialism and Other Essays (pp. 53–74). New York: Oxford UP.

Stubenberg, L. (2016). Neutral Monism. In Stanford Encyclopedia of Philosophy. Retrieved from https://plato.stanford.edu/entries/neutral-monism/

Torre, J. B., & Lieberman, M. D. (2018). Putting feelings into words: Affect labeling as implicit emotion regulation. Emotion Review, 10(2), 116–124.

van Fraassen, B. (1980). The Scientific Image. Oxford: Oxford UP.

Worrall, J. (1989). Structural realism: The best of both worlds? Dialectica, 43, 99–124.

